# Wee1 inhibition decouples Cdk1 and Plk1 activities: a role for gradual Cdk1 activation throughout G2 phase

**DOI:** 10.1101/2024.03.15.585186

**Authors:** Karen Akopyan, Arne Lindqvist

## Abstract

At completion of DNA replication, the mitotic kinases CDK1 and PLK1 are activated. Their activities increase slowly through early G2 phase, but the reason for this low-level activity before mitotic entry is not clear. Using a combination of experiments and mathematical modelling, we find that gradual CDK1 activation through G2 phase stimulates production of mitotic factors and coordinates activation of CDK1 and PLK1. We find that inhibition of CDK1 during G2 phase limits transcription of mitotic factors.

Conversely, the duration of premature mitosis by forced activation of CDK1 is inversely related to the time-point in G2 when mitosis is triggered. Forced CDK1 activation not only leads to a lack of mitotic factors, but also decouples CDK1 and PLK1 activation. Accordingly, we find that duration of forced mitosis by WEE1 inhibition can be partially rescued by expression of constitutively active PLK1. Our results show a function for slow mitotic kinase activation through G2 phase and suggests a mechanism for how the timing of mitotic entry is linked to preparation for mitosis.

**Highlights:**

- Slow Cdk1 activation through G2 phase coordinates expression of mitotic proteins and activation of mitotic kinases
- The duration of forced mitosis by WEE1 inhibition depends on when in G2 mitosis is triggered
- Forced CDK1 activation decouples CDK1 and PLK1 activation
- The duration of forced mitosis can be partially rescued by expression of constitutively active PLK1

## Introduction

For most mammalian cells, there is a long pause between DNA replication and cell division that is referred to as G2 phase. Why G2 phase frequently takes hours is not clear. Initial observations led to the idea that G2 phase was required for protein synthesis (Donnelly and Sisken, 1967), among other of Cyclins that helps trigger mitosis (Minshull et al., 1989). However, more recent work showed that protein synthesis in G2 phase is not strictly required for mitotic entry (Lockhead et al., 2020). Moreover, inhibition of Wee1 can force mitotic entry even before a cell has entered G2 phase, raising the question why a long G2 phase exists (Aarts et al., 2012).

Mitosis is triggered by Cdk1 in complex with Cyclin A or Cyclin B. This Cdk1 activity is suppressed by ongoing DNA replication, ensuring that mitosis is not triggered during S phase (Lemmens et al., 2018; Saldivar et al., 2018). Once DNA replication is complete the suppression is lifted, and initial Cdk1 activity can be detected at the S/G2 border. This low level Cdk1 activity gradually increases throughout G2 phase, until increasing in an exponential fashion which eventually triggers mitosis (Akopyan et al., 2014).

The increase of Cdk1 activity throughout G2 phase can be attributed to the presence of multiple feedback loops (Lindqvist et al., 2009). First, Cdk1 activity stimulates transcription of factors that will further increase Cdk1 activation (Laoukili et al., 2008; Major et al., 2004; Saldivar et al., 2018; Wierstra and Alves, 2006). Second, Cdk1 activity posttranslationally modifies regulators of Cdk1, thereby increasing Cdk1 activity. Whereas some feedback loops are direct, such as activation of Cdc25s (Hoffmann et al., 1993), other involve multiple components such as regulation of Wee1 by Plk1 (Watanabe et al., 2005), which in turn is activated after Cdk1-mediated phosphorylation of the Aurora A cofactor Bora (Tavernier et al., 2015; Thomas et al., 2016; Vigneron et al., 2018). Due to the feedback loops, Cdk1 has been shown to function as a bistable switch (Lindqvist et al., 2007; Pomerening et al., 2003; Rata et al., 2018; Sha et al., 2003).

Importantly, the presence of bistability shows that once initiated, Cdk1 activity will progress until reaching an active steady state but does not say how long it will take for the switch to flip. Apparently, for most human cells the duration is several hours, but what possible benefits could be for a long G2 phase are unknown.

Here we have investigated the feedback that stimulates Cdk1 activation. By combining experiments with a data-driven mathematical model, we find that slow Cdk1 activation through G2 phase coordinates expression of proteins required for mitosis with activation of Plk1. Correspondingly, we find that the duration of a forced mitosis after Wee1 inhibition is inversely related to the time-point in G2 phase when mitosis is triggered. Further, we find that the duration of forced mitosis can be partially rescued by expression of active Plk1.

## Results

### 1. Cdk1 regulates Cyclin B1 accumulation during G2 phase

To test the contribution of cell-cycle dependent kinases on production of mitotic proteins, we sought to monitor protein levels in live cells. We reasoned that Cyclin B1 could function as a proxy for a regulated mitotic protein, as its expression is undetectable in G1 phase, remains limited during early S phase, and increases dramatically during G2 phase (Pines and Hunter, 1989). We therefore utilized a cell line in which we have targeted endogenous Cyclin B1 with YFP (Akopyan et al., 2014). We monitored single U2OS Cyclin B1-YFP cells growing on micropatterns to decrease variability due to cell-extrinsic factors and to increase quantification accuracy of Cyclin B1-YFP levels over time (Akopyan et al., 2014) (**Fig 1A**).

**Figure 1.**
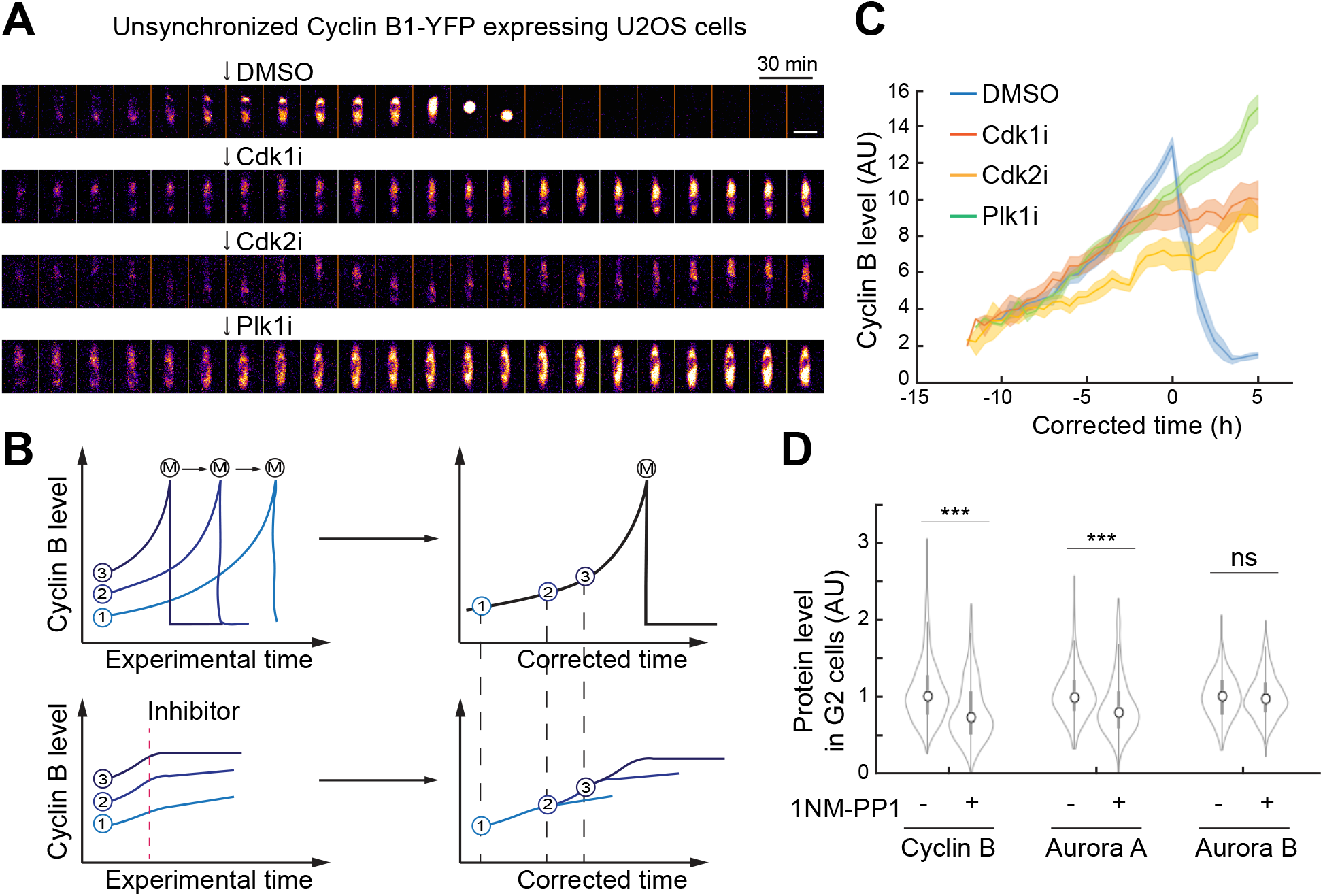
Cdk1 regulates Cyclin B accumulation in G2 phase. **A**. Example of unsynchronized U2OS Cyclin B1-YFP cells growing on micropatterns. Lighter colors indicate higher Cyclin B1-YFP fluorescence. Arrows indicate addition of DMSO or inhibitors. Time lapse 30 min. Scale bar 20 μm. **B**. Schematic of the *in-silico* synchronization setup. DMSO treated cells are synchronized in mitosis (top). The resulting Cyclin B1-YFP fluorescence curve (top right) is used to fit Cyclin B1-YFP fluorescence of individual cells before kinase inhibitor treatment (bottom). **C**. Quantification of Cyclin B1-YFP fluorescence of cells treated with DMSO or indicated kinase inhibitors after in-silico synchronization as in B. Graph shows average and standard error of Cyclin B1-YFP fluorescence. Please note that only Cyclin B1-YFP fluorescence after kinase inhibitor addition is plotted. **D**. Quantification of Cyclin B1, Aurora A and Aurora B immunofluorescence in 4N U2OS-Cdk1as cells after 2h treatment with 1NM-PP1.

We next added small molecule kinase inhibitors to unsynchronized cells and quantified Cyclin B1-YFP levels over time. We chose to test inhibitors of Cdk1 (RO3306), Cdk2 (NU6140), and Plk1 (BI2536), three kinases that are central for cell cycle progression and active throughout G2 phase (**Fig 1A**). To pinpoint if and when an inhibitor impacted Cyclin B1-YFP accumulation, we devised an *in-silico* synchronization scheme. We first aligned control cells on mitotic entry, creating a trendline of Cyclin B1-YFP accumulation over time (**Fig 1B**, top panels, and **Fig1C**, DMSO). Using the Cyclin B1-YFP levels before inhibitor addition, we next aligned all cells to the trendline, thereby assigning an approximate cell cycle position for each cell (**Fig 1B**, lower panel). We then plotted Cyclin B1-YFP accumulation after addition of each inhibitor and compared it to the trendline of control cells. Importantly, this approach is designed to identify if and when in the cell cycle Cyclin B1-YFP accumulation is initially affected by inhibitor addition but may not be quantitative for the magnitude of changes after an effect is observed.

Addition of NU6140 led to decreased Cyclin B1-YFP accumulation more than 6h before mitosis, whereas addition of RO3306 or BI2536 led to decreased Cyclin B1-YFP accumulation approximately 3h before mitosis (**Fig 1C**). The data is in agreement with the activity profiles of the targets of the inhibitors, as Cdk2 activity is progressively increasing through S phase, and Cdk1 and Plk1 are progressively increasing through G2 phase (Akopyan et al., 2014; Spencer et al., 2013). Our results would be consistent with a model in which Cdk2 ensures a base-line of Cyclin B1 accumulation already from S-phase, whereas Cdk1 activity and to some extent Plk1 activity enables the increase in Cyclin B1 levels during G2 phase.

To test whether Cdk1 activity drives the increase of Cyclin B1 levels in G2 phase, we used immunofluorescence to quantify endogenous Cyclin B1 levels in an unsynchronized population of U2OS Cdk1as cells, allowing selective inhibition of Cdk1 (Rata et al., 2018). Focusing on cells with a 4N DNA content, we note that cells treated for 2 hours with the bulky ATP analogue 1NM-PP1 show reduced staining for Cyclin B1 (**Fig 1D**). We conclude that Cdk1 activity increases Cyclin B1 levels in G2 phase.

### 2. Cdk1 stimulates synthesis of mitotic factors and of proteins that boost its own activity

The increase in Cyclin B1 levels late in the cell cycle largely stems from upregulated transcription, and Cdk activity has been implicated in increasing transcription of Cyclin B1, in particular by modification of FoxM1 (Laoukili et al., 2008; Lüscher-Firzlaff et al., 2006; Major et al., 2004; Saldivar et al., 2018). To test if Cdk1 inhibition affects Cyclin B1 production, we treated unsynchronized U2OS cells with the translation inhibitor cycloheximide (CHX) and analyzed Cyclin B1 content in G2 cells by quantitative immunofluorescence. Whereas RO3306 addition for 2 hours reduced Cyclin B1 levels over control cells, treatment with CHX or CHX together with RO3306 showed a similar decline in Cyclin B1 levels (**Fig 2A**), suggesting that RO3306 addition can affect production of Cyclin B1.

**Figure 2.**
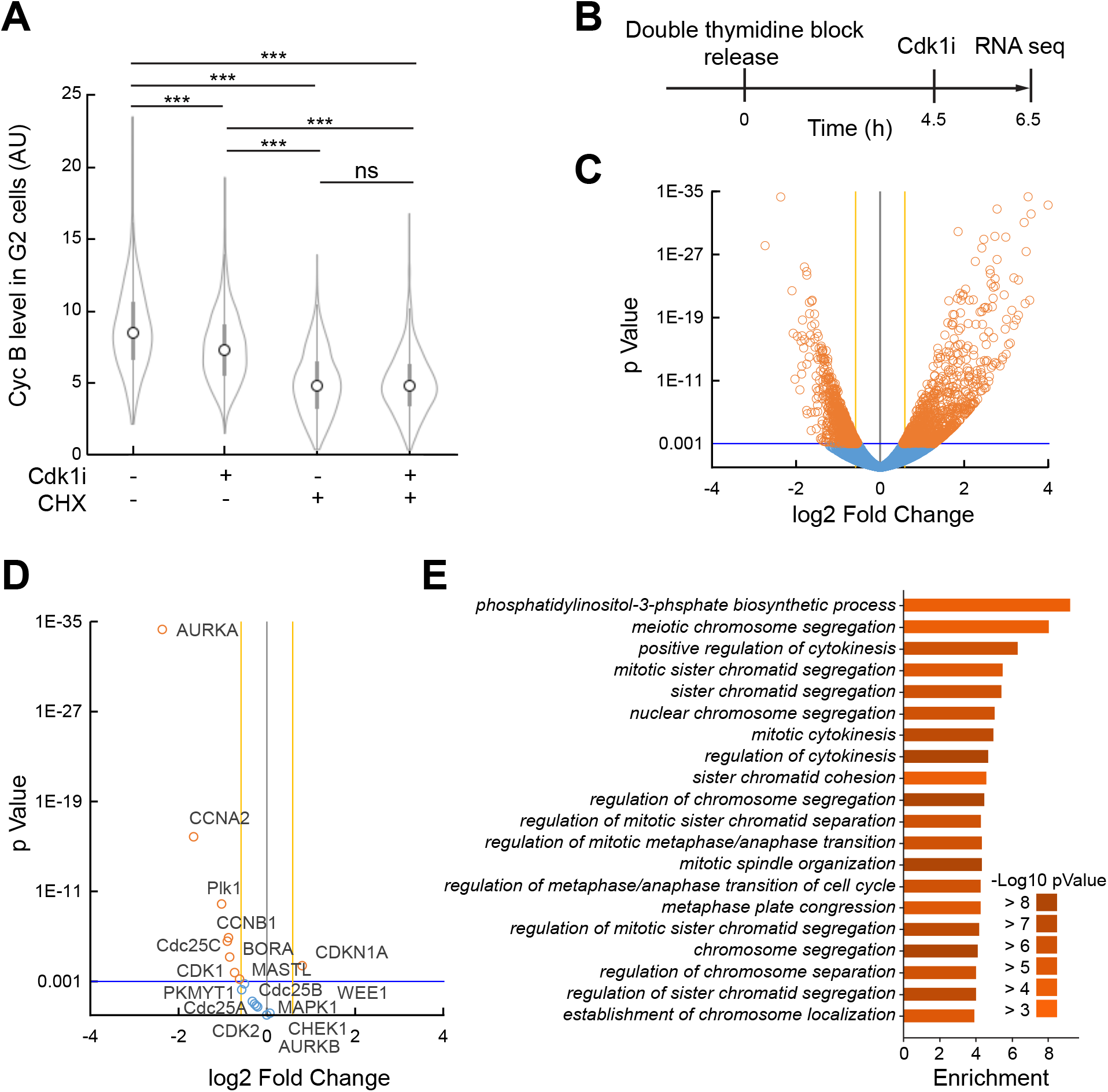
Cdk1 regulates transcription of mitotic factors. **A**. Quantification of Cyclin B1 immunofluorescence in 4N U2OS cells treated with Cdk1 inhibitor (RO3306), cycloheximide or both for 2 hours. The G2 population was separated *in silico* based on DAPI. **B**. Schematic of setup for RNA sequencing. HeLa cells were released after double thymidine synchronization. After 4.5 hours Cdk1 inhibitor (RO3306) or DMSO was added. After 2 hours cells were harvested for RNAseq analysis. **C, D**. Volcano plots show log2 fold change between treated (RO3306) and non-treated (DMSO) normalized gene expressions (x-axis), plotted versus the p-value (y-axis). Orange circles represent differentially expressed genes and blue circles represent genes with similar expression. **E**. Gene Ontology and p-values based on C.

To test if RO3306 addition affected transcription of Cyclin B1, we next synchronized HeLa cells (**Fig 2B**) and sequenced poly A-enriched RNA. In line with the previous results, we find Cyclin B1 mRNA levels to be reduced after RO3306 addition in G2 phase (**Fig 2C, D)**. The largest decrease in mRNA levels after RO-3306 treatment in G2 phase was for the mitotic kinase Aurora A. We verified that Aurora A expression was affected by addition of 1NMPP1 to CDK1as cells in G2 phase, whereas the related mitotic kinase Aurora B was not affected at mRNA level after RO3306 addition or at protein level after 1NMPP1 addition (**Fig 1D, 2D)**. Thus, at least for Cyclin B1, Aurora A, and Aurora B, similar trends are seen on mRNA and protein levels using two different approaches of CDK1 inhibition in G2 phase.

Except Cyclin B1 and Aurora A, the mRNA levels of many genes are affected by RO-3306 treatment in G2 phase (**Fig 2 C, D)**. Using Gene Ontology (GO) analysis, we find an enrichment for various processes that are important for mitosis, including mitotic spindle formation, regulation of chromosome segregation and cytokinesis. This suggests that Cdk1 activity in G2 phase can stimulate the production of multiple proteins that regulate mitosis (**Fig 2E**).

Cdk activity is increasing in an exponential fashion during G2 phase, which is attributed to the presence of multiple feedback loops. These feedback loops can be divided into direct feedback on cyclin-Cdk complex levels, inner feedback loops that directly activates Cyclin-Cdk, and outer feedback loops that indirectly activate Cyclin-Cdk (Lindqvist et al., 2009). How large part transcriptional regulation plays for the various feedback systems is not clear. We therefore searched for changes in mRNA levels that would be expected to impact on Cdk activity itself. Apart from Cyclin B1, Cyclin A2 and Cdk1 are among the mRNAs that decreased upon RO3306 addition in G2 phase (**Fig 2D**). This suggests that Cdk1 activation in G2 phase stimulates the presence of more Cyclin-Cdk1 complexes. However, with the exception of Cdc25C, mRNA levels for the direct Cyclin-Cdk regulators Cdc25A, Cdc25B, Wee1 and Myt1 were unaffected by RO3306 addition, suggesting that transcriptional regulation is limited for the inner feedback loops that activate Cdk1 through post-translational modifications. Interestingly, mRNA for Aurora A, Bora, and Plk1, which indirectly can amplify Cdk1 activity, was reduced after RO3306 addition in G2 phase. Thus, transcriptional regulation seems to exist within the outer feedback loops for Cdk1 activation.

In conclusion, our data would be consistent with a model in which Cdk1 activity in G2 phase stimulates both the transcription of many proteins with mitotic functions and of proteins that stimulate further Cdk1 activation. The transcriptional component seems to be strong in particular for the kinases Aurora A and Plk1 that in addition to stimulate Cdk1 activity also are key regulators of mitotic processes.

### 3. Model

To better understand the implications of Cdk-mediated regulatory feedback, we decided to use a simulation approach based on experimental data. We sought to create a model that balances overparameterization when fitting to data by only including a limited set of main players that regulate Cdk activity. At the same time, we sought to include an updated framework of cell cycle regulation concerning the large part played by DNA replication on regulating interphase (Lemmens and Lindqvist, 2019) (**Fig 3A**).

**Figure 3.**
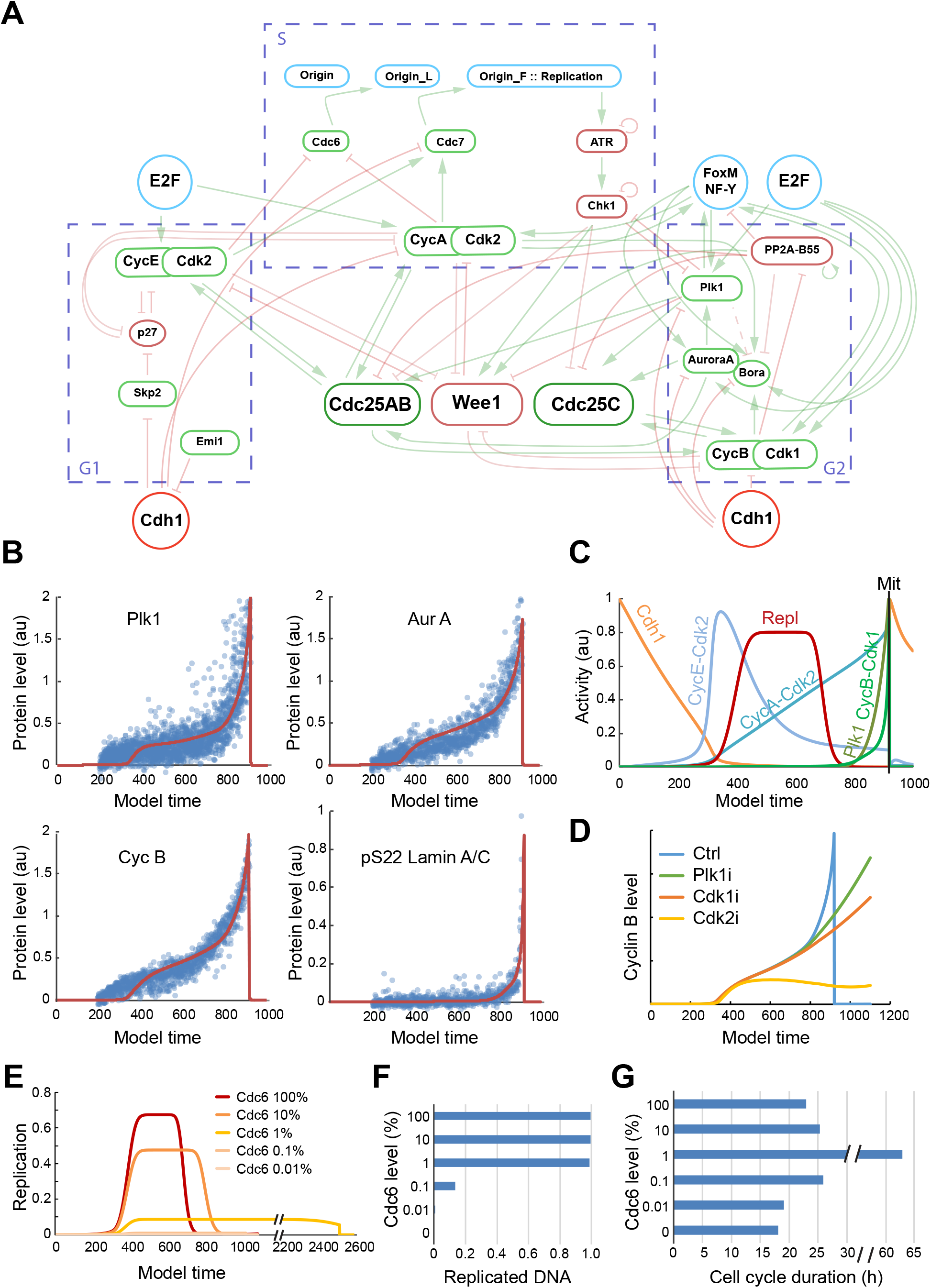
Mathematical model of the cell cycle. **A**. Schematic representation of the mathematical model. **B**. Model prediction (red line) of the protein level dynamics after parameter estimation based on quantitative immunofluorescence of indicated proteins in U2OS cells from (Akopyan et al., 2014) (blue dots). **C**. Model estimation of selected cell cycle activities. **D**. Model prediction of Cyclin B level dynamics after inhibition of Cdk1, Cdk2 or Plk1. **E, F, G**. Simulation of the replication dynamics, total replicated DNA and cell cycle duration at different levels of Cdc6. 100% Cdc6 refers to Cdc6 level used in C.

To start the cell cycle, we used a mathematical model of the G1/S transition from Novák and coauthors, which has been fitted to experimental data (Barr et al., 2016). In S phase, Cdk2 complexes inactivate origin licensing (in the model for simplicity simulated as Cdc6) and activate origin firing (simulated as Cdc7), which initiates DNA replication. While DNA replication is ongoing it inhibits (simulated as ATR and Chk1) the activities of main cell cycle kinases. When DNA replication is completed in G2 phase, various feedback loops in the mitotic entry network ensures that Plk1 and Cdk1 activities will rise and bring the cell to mitosis. The transcriptional feedback through Cdk1 based on **Figure 2** is simulated as FoxM1. For simplicity, both Cyclin A-Cdk1 and Cyclin B-Cdk1 are simulated as Cyclin B-Cdk1. For a detailed description of the model, see **Appendix 1**.

After an initial manual selection of parameters, we used experimental data from our previous work (Akopyan et al., 2014) to fit model parameters to reflect changes in protein levels of Cyclin B, Plk1 and Aurora A. Similarly, we fitted model parameters to reflect Cdk1 activity based on the level of phosphorylated Lamin A/C (**Fig 3B**). The model containing updated parameters is in agreement with main trends of order and shape of main cell cycle regulators and their activities (**Fig 3C**).

To test the accuracy of the model, we decided to reproduce the situation described in **Figure 1C**. We therefore simulated inhibition of Cdk1, Cdk2 or Plk1 and monitored the resulting Cyclin B level. The simulation recapitulated the trends of the experimental data, indicating that the model can describe main aspects of how Cdk1, Cdk2, and Plk1 are linked to Cyclin B production (**Fig 1C and 3D**).

We next sought to test the predictive power of the model for a variable that was not fitted to experimental data. A key aspect of the model is that DNA replication functions as a signaling component that limits G2 specific activities and thereby coordinate S phase and mitosis (Lemmens and Lindqvist, 2019). Simulating a reduction in DNA replication origin licensing by reducing the levels of Cdc6, we see that DNA replication occurs at a lower speed, and consequently, takes longer (**Fig 3E**). As the levels of Cdc6 decrease, the inhibition from DNA replication on mitotic kinases is not sustained until completion of DNA replication, and the cell eventually enters mitosis without replicating all DNA (**Fig 3F**).

Importantly, whereas the cell cycle is prolonged at intermediate levels of Cdc6, very low levels of Cdc6 leads to premature activation of mitotic kinases and consequently, to a shorter cell cycle (**Fig 3G**). These data are in agreement with observations using degron-tagged Cdc6 in cells (Lemmens et al., 2018), indicating that the model can recapitulate the basic logic of how DNA replication coordinates activities through S and G2 phases.

### 4. Mitotic duration after Wee1 inhibition depends on when in G2 phase Wee1 inhibitors are added

Having established that the model can recapitulate main aspects of how Cdk1, Cdk2, and Plk1 activities and associated protein levels rise through S and G2 phase, we next focused on the importance of Cdk1-dependent feedback. We therefore removed Wee1 activity *in silico* at various points in G2 phase. Not surprisingly, the model rapidly gained full CDK1 activity and entered mitosis (**Fig 4A**). When Wee1 activity was removed in early G2 phase there was a small delay before mitosis, in which G2 transcription was active while CDK1 activity was rising (**Fig 4A and B**). This delay ensured that when Wee1 activity was removed, mitosis only occurred once a threshold level of Cyclin B1 was reached, which corresponded to the Cyclin B level present in mid-G2 phase (**Fig 4C**).

**Figure 4.**
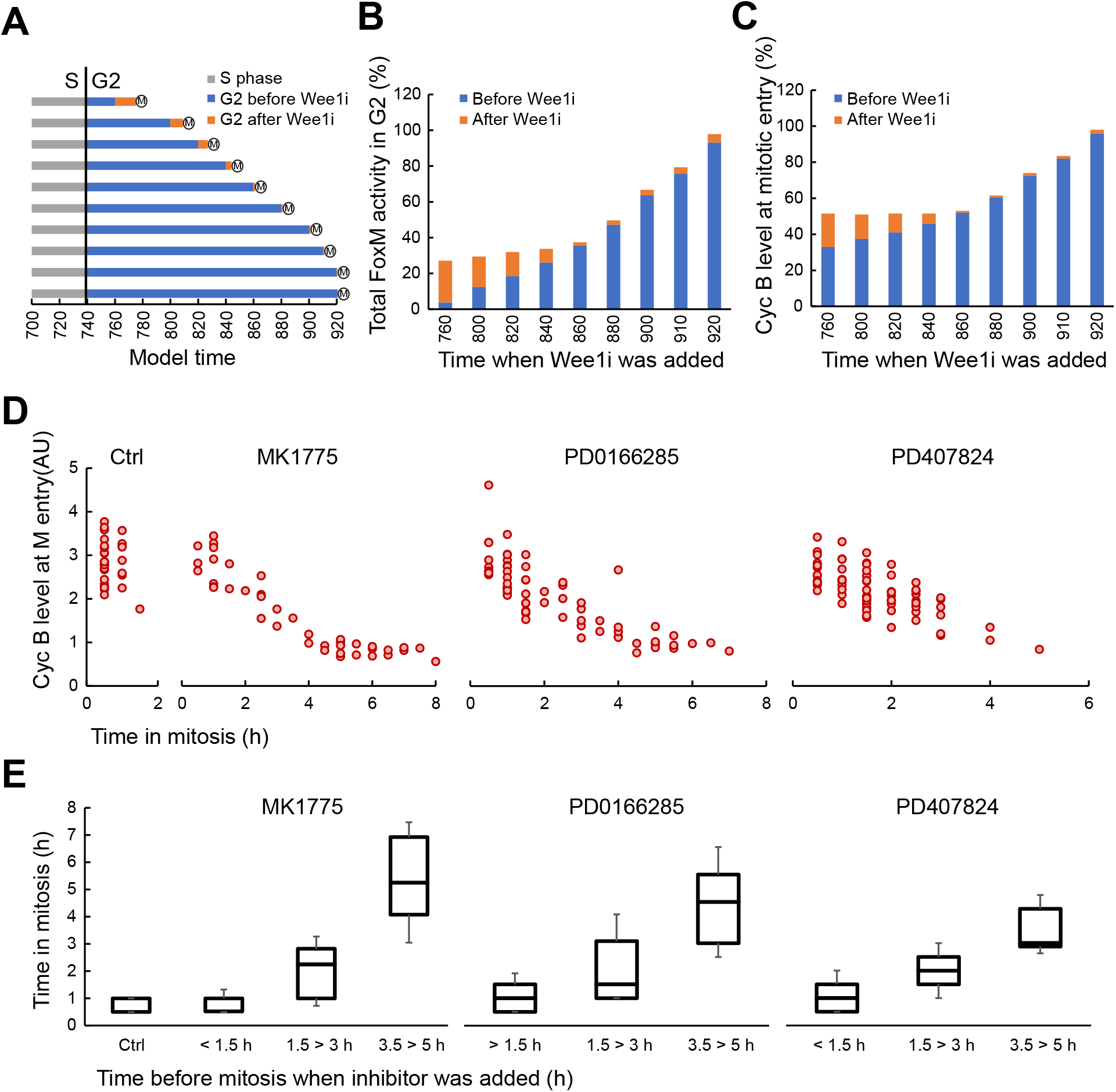
Mitotic duration after Wee1 inhibition depends on when in G2 phase Wee1 inhibitors are added. **A**. Model prediction of G2 duration after Wee1 inhibition at different timepoints in G2 phase. **B**. Model prediction of accumulated FoxM activity at mitotic entry after Wee1 inhibition at different timepoints in G2 phase. 100% denotes mitotic levels in absence of Wee1 inhibition. **C**. Model prediction of Cyclin B level at mitotic entry after Wee1 inhibition at different timepoints in G2 phase. 100% denotes mitotic levels in absence of Wee1 inhibition. **D, E**. U2OS Cyclin B1-YFP cells were monitored by time-lapse microscopy upon addition of Wee1 inhibitors. Three different inhibitors were used - MK1775 (1 μM), PD0166285 (1 μM) and PD407824 (5 μM). D, duration of mitosis (x-axis) is plotted versus Cyclin B1-YFP level at mitotic entry (y-axis). **E**. Duration of mitosis (y-axis) is plotted versus estimated time before mitosis should Wee1 inhibitors not have been added (x-axis). The estimate is based on Cyclin B1-YFP accumulation of control cells in the same experiment (Fig S2).

We next tested this prediction experimentally by addition of three different Wee1 inhibitors (MK1775, PD0166285 and PD407824) to cells in which Cyclin B1 is tagged to YFP in the endogenous locus. As expected, forcing premature mitosis by Wee1 inhibitor addition led to the presence of mitotic cells with reduced levels of Cyclin B1-YFP (**Fig 4D and S1)**. In accordance with simulation, all mitotic cells contained a basal expression of Cyclin B1-YFP. However, the detected minimal level of Cyclin B1-YFP in mitosis is lower than minimal Cyclin B during simulation, possibly reflecting that both Cyclin A2-Cdk1 and Cyclin B1-Cdk1 are grouped as Cyclin B-Cdk1 during simulation (**Fig 4C and D)**.

To study the consequences of Wee1 inhibition, we next monitored mitotic progression of single Wee1 inhibitor treated cells. We find that cells that passed mitosis with similar timings as control cells also contained Cyclin B-YFP levels similar to control cells. In contrast, cells that showed a prolonged mitosis contained lower levels of Cyclin B-YFP. The level of Cyclin B-YFP before Wee1 inhibitor addition was inversely related to the duration of mitosis (**Fig 4D and S1**).

Using an *in silico* approach, we estimated how far into G2 phase each cell was at the time of Wee1 inhibition. Similar to in **Figure 1B**, the estimate is based on relating the Cyclin B-YFP levels of each cell to the average trendline of how Cyclin B-YFP levels increase throughout G2 phase (**Fig S2**). We find that mitotic duration is inversely related to when in G2 phase Wee1 inhibitors are added (**Fig 4E**).

Thus, our findings indicate that forced Cdk activation in G2 phase results in mitotic entry before all mitotic proteins have accumulated to levels observed in unperturbed cells. We find that Wee1 inhibition is not toxic for mitotic progression if added in late G2 phase. However, if the inhibitor is added earlier in G2 phase, it leads to mitotic delays. Taken together, this indicates that G2 progression and intact Cdk regulation during G2 phase is important for mitotic progression.

### 5. Decoupled Cdk1 and Plk1 activities contribute to prolonged mitosis after Wee1 inhibition

Apart from production of mitotic proteins, Cdk dependent feedback involves activation of the mitotic kinase Plk1 that is crucial for mitotic progression (Steegmaier et al., 2007). We therefore simulated how Plk1 activity would be affected by removal of Wee1 activity at various points in G2 phase. In the model, we find that the Aurora A/Bora-dependent feedback loops that stimulate Plk1 activity depends heavily on gradual activation of Cdk1 (**Fig 5A)**. According to simulations, if Wee1i is added early in G2 phase the cell rapidly develops high Cdk1 activity, intermediate levels of Cyclin B (which in the model functions as a proxy for a Cdk1-dependent transcriptional target), and low Plk1 activity. In contrast, if Wee1i is added late in G2 phase, this affect is less apparent as a large part of both Cdk1-dependent transcription and Plk1 activation already occurred earlier in G2 phase (**Fig 5B)**.

**Figure 5.**
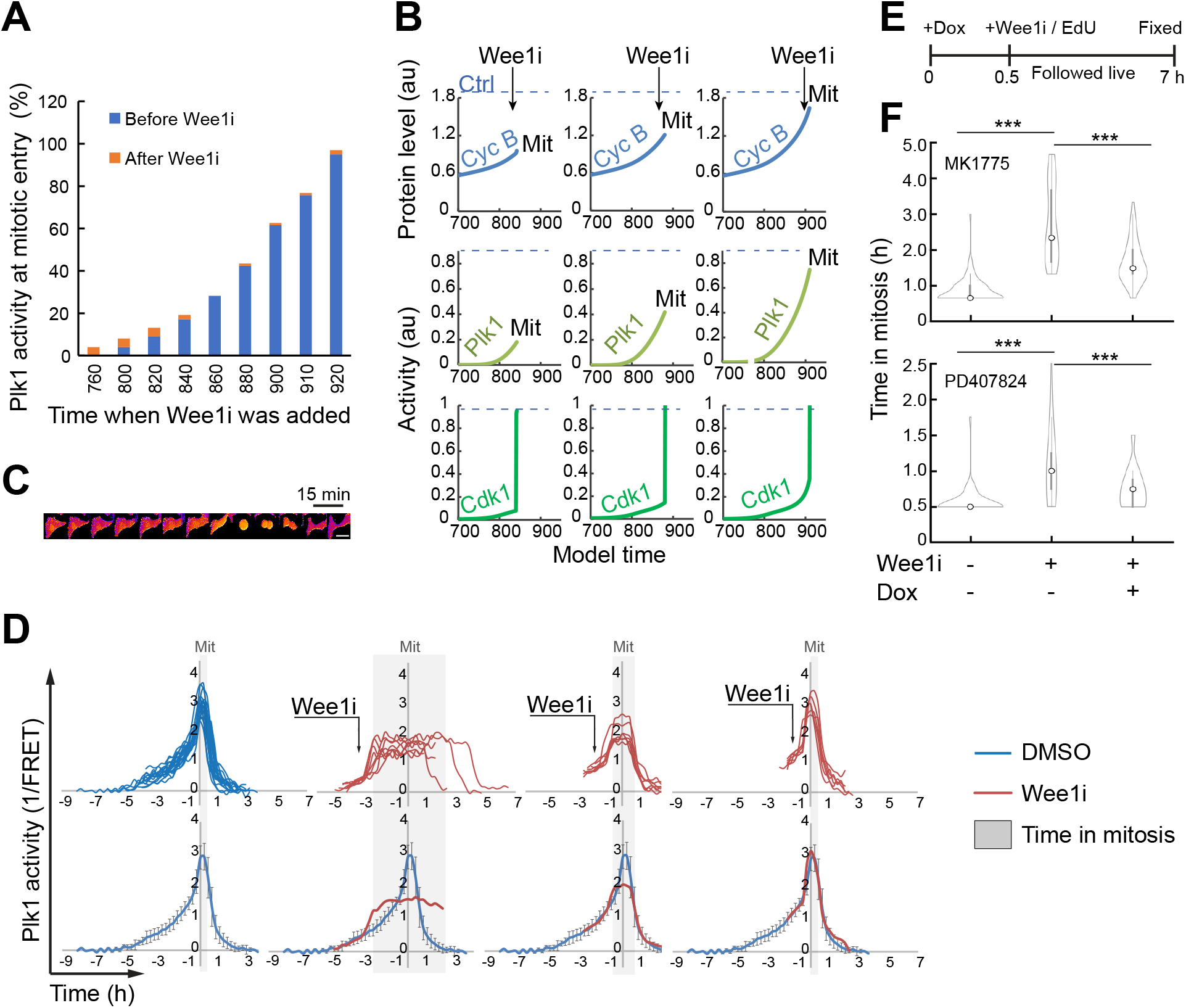
Decoupled Cdk1 and Plk1 activities contribute to prolonged mitosis after Wee1 inhibition. **A**. Model prediction of Plk1 activity at mitotic entry after Wee1 inhibition at different timepoints in G2 phase. 100% denotes mitotic levels in absence of Wee1 inhibition. **B**. Model prediction of Cyclin B level (top) Plk1 activity (middle) and Cdk1 activity (bottom) after inhibition of Wee1 at different time points in G2 phase. The dotted line in each graph represents the levels at mitotic entry when Wee1 is not artificially inhibited. Mit indicates mitosis. **C**. Time lapse imaging of U2OS cell expressing Plk1-FRET probe. Lighter colors indicate phosphorylated probe. Time lapse 15 min. Scale bar 20 μm. **D**. Dynamics of Plk1 activity in single U2OS cells expressing Plk1-FRET probe (top) and average with standard deviation (bottom). Blue line in bottom panel corresponds to average values of cells treated with DMSO only and red line to average for cells treated with Wee1i (MK1775). Arrows represent the time when MK1775 was added. The grey area indicates the duration of mitosis. **E**. Schematic of the setup for **F.** **F**. Mitotic duration of U2TR - Plk1-T210D cells treated with DMSO (Ctrl), 1 μM MK1775 or 3 μM PD407824 (Wee1i) and Doxycycline (Dox) as indicated. Cells that entered mitosis between 80 and 120 minutes after Wee1i addition were followed to ensure comparable populations and to exclude cells that were in late G2 phase upon Wee1i addition. Cells were monitored after fixation and only EdU negative cells (indicating that they were not in S phase upon inhibitor addition) were included in the analysis.

To test this prediction experimentally we used U2OS cells expressing a FRET-based probe that monitors Plk1 target phosphorylation in single cells **(Fig 5C)**. We monitored FRET while adding Wee1 inhibitor and followed cells through mitosis. Based on the FRET signal of control cells, we estimated the time remaining to mitosis should no inhibitor have been added **(Fig 5D, left)**. Similar to the model, we find that cells entering mitosis after Wee1 inhibitor addition in early G2 phase were entering mitosis with low Plk1 activity **(Fig 5D, second left)**. In contrast, when Wee1 inhibitors were added in late G2 phase, we detect no difference in mitotic Plk1 activity compared to control cells **(Fig 5D, right)**. Similar to Cyclin B-YFP levels, the duration of mitosis inversely correlates with the level of Plk1 activity in mitotic entry (**Fig 5D, grey rectangles)**.

To test whether deficient Plk1 activation after Wee1i addition in G2 phase contributes to prolonged mitosis, we sought to increase Plk1 activity after Wee1i was added. To this end, we used U2TR - Plk1-T210D cells, in which a constitutively active form of Plk1 can by expressed by addition of Doxycycline. As induction of expression of Plk1 T210D requires both transcription and translation, we added Doxocycline 30 min before Wee1i to minimize Plk1 T210D expression before Wee1 inhibition (**Fig 5E)**. As expected, the mitotic duration increased after treating G2 cells with Wee1i. This increase was partially rescued by expression of Plk1 T210D, suggesting that part of the mitotic delay after Wee1 inhibition in G2 phase depends on a lack of active Plk1 **(Fig 5F)**.

## Discussion

“In G2 phase, a cell prepares for mitosis” is a common statement during undergraduate education, but how this preparation is controlled is less clear. Here we find that gradual Cdk1 activation through G2 phase both ensures production of mitotic proteins and coordinates activation of mitotic kinases. Using a combination of simulation and experiments, we find that Cdk1 involvement in feedback loops underlies this coordination.

It is now one decade since we reported that Cdk1 and Plk1 are activated upon completion of DNA replication, but the reason for a gradual activation initiated several hours before mitosis has been enigmatic (Akopyan et al., 2014). Full Cdk1 activation results in mitotic entry, but before such an activity is reached, a gradual activation takes place throughout G2 phase. As Cdk1 activation depends on accumulation of Cyclins, the coupling of production of both Cyclins and other mitotic regulators to Cdk1 activity ensures the presence of proteins required for mitosis when a sufficient pool of Cyclin-Cdk1 is present.

Whereas the transcriptional regulation ensures the presence of proteins, their activities are regulated by posttranslational modifications. We find that the gradual activation of Cdk1 ensures that both Cdk1 and Plk1 are active upon mitotic entry. Thus, a gradual activation throughout G2 phase allows coordinated activities of Cdk1 and Plk1 when mitosis occurs. We find that this coordinated accumulation of mitotic factors and Cdk1 and Plk1 activities are disrupted when Cdk1-dependent feedback loops are shortcut by addition of Wee1 inhibitors. The resulting increased mitotic duration is likely also affected by slower activation kinetics as previously reported (Araujo et al., 2016).

Wee1 inhibitors are used in clinical trials with the rationales that Wee1 inhibition can overrule a G2 checkpoint and that cells are forced to mitosis prematurely which leads to mitotic delays and mitotic catastrophe (Kong and Mehanna, 2021). The inhibitor AZD1775 has been suggested to also target Plk1, although whether Plk1 inhibition occurs at clinically relevant concentrations has been questioned (Serpico et al., 2019; Wright et al., 2017). Here, we find that the magnitude of a mitotic delay depends on the timepoint in G2 phase when Wee1 inhibitors are added. When Wee1 inhibitors were added close to mitosis, only minimal effects on mitotic duration were observed. This is in stark contrast to Plk1 inhibitors, for which a mitotic delay is observed even if added close to mitosis (Steegmaier et al., 2007). Further, measurement of Plk1 activity in single cells using a FRET-based reporter show no decrease in Plk1 activity upon addition of a Wee1 inhibitor. Rather, as predicted by modelling, cells retained similar Plk1 activity as they had in G2 phase at the time Wee1 inhibitor was added (**Fig 5**).

Our findings show a mechanistic rationale for why both Wee1 inhibitor treatment and genetic abrogation of Wee1 shows synergistic effects with Plk1 inhibition (Wright et al., 2017). Further, they are consistent with synergistic effects between Aurora A, the upstream kinase activating Plk1, and Wee1 inhibition in head and neck cancer models (Lee et al., 2019). To date, 79 clinical trials using Wee1 inhibitors are reported, and a number of suggested biomarkers for treatment efficiency is available (Wang et al., 2024). Our finding that expression of active Plk1 partially can rescue prolonged mitosis after Wee1 inhibition opens for further investigations on whether combined Wee1 and Plk1 inhibition would be beneficial and whether Plk1 activity can predict Wee1 inhibitor treatment efficiency.

## Materials and Methods

### Cells and Cell culture

U2OS CyclinB1-YFP, and hTERT RPE1 Cyclin B1-YFP cells were from (Akopyan et al., 2014). U2OS, U2OS Plk1-FRET and U2TR - Plk1-T210D cells were from (Macůrek et al., 2008). U2OS-Cdk1as cells were from (Rata et al., 2018).

U2OS and HeLa cells were cultured in DMEM plus GlutaMAX (Invitrogen) supplemented with either 6% (for U2OS and U2OS CyclinB1-YFP) or 10% (for HeLa, U2OS-Cdk1as, U2OS Plk1-FRET and U2TR - Plk1-T210D), heat-inactivated FBS (HyClone) and 1% Penicillin-Streptomycin (HyClone). hTERT RPE1 Cyclin B1-YFP were cultured in DMEM/F12 plus GlutaMAX (Invitrogen) supplemented with 10% heat-inactivated FBS (HyClone) and 1% Penicillin-Streptomycin (HyClone). All cells were maintained in an incubator with controlled conditions at 37°C and 5% CO_2_.

For live-cell imaging experiments, the medium of the cells was changed 24 hr prior to imaging to CO_2_-independent medium (Leibovitz 15; Invitrogen) supplemented with 6% or 10% heat-inactivated FBS and 1% Penicillin-Streptomycin (HyClone).

### Cell synchronization

Cells were synchronized for 20 h with 2.5 mM thymidine (Sigma-Aldrich) and released into fresh medium for 12 h, followed by 2.5 mM thymidine for an additional 14 h.

### Live-Cell Imaging

A DeltaVision Spectris Imaging System with a 20× air objective (NA 0.75 ) or a Leica DMI6000 Imaging System with a 40× air objective (NA 0.85) or 20× air objective (NA 0.40) were used to follow the cells live. Images were acquired with an interval of 15–30 min.

The images were analyzed using custom ImageJ (https://imagej.nih.gov/ij/) or MATLAB scripts.

Imaging and quantification of the CFP/YFP emission ratio FRET cells was performed as described in (Hukasova et al., 2012).

Fibronectin-coated micropatterns (CYTOO) were used as described in (Akopyan et al., 2014).

### Fixed-cell imaging

Cells were seeded in a 96-well imaging plate (BD Falcon) 24 h prior fixation. The fixation (3.7% formaldehyde (Sigma Aldrich) for 5 min) was followed by permeabilization for 2 min in ice-cold methanol. Fixed cells were incubated in blocking solution: TBS-T (TBS with 0.1% Tween20) supplemented with 2% bovine albumin serum for 1 h at room temperature. Then cells were incubated in in blocking solution with primary antibodies overnight at 4°C, washed 3 times in TBS-T and incubated with secondary antibodies and DAPI for 1h at room temperature. Samples were washed 3 times in TBS-T and stored in TBS or PBS.

EdU-Click chemistry was performed by incubation in Tris (pH 6.8) – 100mM, CuSO4 – 1mM, Ascorbic acid– 10mM and Azid Flourophore (#A10277, Invitrogen) – 3μM for 30 min at room temperature.

Images were acquired using an ImageXpress microscope with 20X objective (NA 0.45) at room temperature. Image analysis and background subtraction was performed as in (Akopyan et al., 2016).

### Antibodies

The following antibodies were used in this study: Cyclin B1 V152 (4135, Cell Signaling), PLK1 (ab14210; Abcam), pTCTP (#5251; Cell Signaling), Aurora A (#4718; Cell Signaling), Aurora B (#3094; Cell Signaling), pLaminA/C (#2026; Cell Signaling), Alexa Fluor 488-Goat anti-Rabbit (#A11008 Life Technologies), Alexa Fluor 555-Goat anti-Mouse (#A21422 Life Technologies).

### Chemicals

The following inhibitors were used in this study:

For Cdk1 inhibition: RO3306 (#217699; CalBiochem) was used at 10 μM; 1NMPP1 (#529581; CalBiochem) 1 μM

For Plk1 inhibition: BI2536 (#S1109; Selleck Chemicals) was used at 100 nM

For Cdk2 inhibition: NU6140 (#238804; CalBiochem) was used at 10 μM

For Wee1 inhibition: MK1775 (Synonyms: AZD1775, Adavosertib, #S1525; Selleckchem) (1 μM), PD0166285 (#S8148; Selleckchem ) (1 μM) and PD407824 (PZ0111; Sigma) (3 or 5 μM).

Doxycycline (D1822; Sigma) 1 μg/ml.

Cycloheximide (C4859; Sigma) 10 μg/ml.

### RNA sequencing

4.5 h after double thymidine block release HeLa cells were treated for 2 h either with Cdk1 inhibitor (RO3306) or DMSO. RNA isolation, sequencing, quality control and mapping of reads were performed by the National Genomics Infrastructure at SciLifeLab. Briefly, strand-specific TruSeq RNA libraries were prepared after poly-A selection and sequenced using an Illumina HiSeq sequencer. Reads were mapped using Tophat, RPKM/FPKM values were calculated using Cufflinks and read counts were calculated using HTSeq.

Data analysis was performed using the DESeq2 package (https://genepattern.github.io/DESeq2) in R.

Gene ontology analysis was performed using <u>G</u>ene <u>O</u>ntology en<u>RI</u>chment ana<u>L</u>ysis and visua<u>L</u>iz<u>A</u>tion tool – Gorilla (https://cbl-gorilla.cs.technion.ac.il)

### Modeling

The mathematical model of the cell cycle was based on differential equations assuming Michaelis– Menten kinetics. A detailed description of all equations and parameters is in Appendix 1.

Solving the equations and parameter estimation were performed using Copasi 4.42, build 284 (https://copasi.org/)

## Acknowledgements

The authors acknowledge support from Science for Life Laboratory, the Knut and Alice Wallenberg Foundation, the National Genomics Infrastructure funded by the Swedish Research Council, and Uppsala Multidisciplinary Center for Advanced Computational Science for assistance with massively parallel sequencing and access to the UPPMAX computational infrastructure. The authors thank Rene Medema, Libor Macurek, and Helfrid Hochegger for reagents.

## REFERENCES

Aarts, M., Sharpe, R., Garcia-Murillas, I., Gevensleben, H., Hurd, M.S., Shumway, S.D., Toniatti, C., Ashworth, A., Turner, N.C., 2012. Forced mitotic entry of S-phase cells as a therapeutic strategy induced by inhibition of WEE1. Cancer Discov. 2, 524–539. 2159-8290.CD-11-0320

Akopyan, K., Lindqvist, A., Müllers, E., 2016. Cell Cycle Dynamics of Proteins and Post-translational Modifications Using Quantitative Immunofluorescence. Methods Mol. Biol. Clifton NJ 1342, 173–183. 10.1007/978-1-4939-2957-3_9

Akopyan, K., Silva Cascales, H., Hukasova, E., Saurin, A.T., Müllers, E., Jaiswal, H., Hollman, D.A.A., Kops, G.J.P.L., Medema, R.H., Lindqvist, A., 2014. Assessing kinetics from fixed cells reveals activation of the mitotic entry network at the S/G2 transition. Mol. Cell 53, 843–853. 10.1016/j.molcel.2014.01.031

Araujo, A.R., Gelens, L., Sheriff, R.S.M., Santos, S.D.M., 2016. Positive Feedback Keeps Duration of Mitosis Temporally Insulated from Upstream Cell-Cycle Events. Mol. Cell 64, 362–375. 10.1016/j.molcel.2016.09.018

Barr, A.R., Heldt, F.S., Zhang, T., Bakal, C., Novák, B., 2016. A Dynamical Framework for the All-or-None G1/S Transition. Cell Syst. 2, 27–37. 10.1016/j.cels.2016.01.001

Bleichert, F., 2019. Mechanisms of replication origin licensing: a structural perspective. Curr. Opin. Struct. Biol., Catalysis and Regulation • Protein Nucleic Interactions 59, 195–204. 10.1016/j.sbi.2019.08.007

Bruinsma, W., Aprelia, M., García-Santisteban, I., Kool, J., Xu, Y.J., Medema, R.H., 2017. Inhibition of Polo-like kinase 1 during the DNA damage response is mediated through loss of Aurora A recruitment by Bora. Oncogene 36, 1840–1848. 10.1038/onc.2016.347

Coulombe, P., Nassar, J., Peiffer, I., Stanojcic, S., Sterkers, Y., Delamarre, A., Bocquet, S., Méchali, M., 2019. The ORC ubiquitin ligase OBI1 promotes DNA replication origin firing. Nat. Commun. 10, 1–14. 10.1038/s41467-019-10321-x

Donnelly, G.M., Sisken, J.E., 1967. RNA and protein synthesis required for entry of cells into mitosis and during the mitotic cycle. Exp. Cell Res. 46, 93–105. 10.1016/0014-4827(67)90412-0

Dutta, A., Bell, S.P., 1997. Initiation of Dna Replication in Eukaryotic Cells. Annu. Rev. Cell Dev. Biol. 13, 293–332. 10.1146/annurev.cellbio.13.1.293

Fattaey, A., Booher, R.N., 1997. Myt1: a Wee1-type kinase that phosphorylates Cdc2 on residue Thr14, in: Meijer, L., Guidet, S., Philippe, M. (Eds.), Progress in Cell Cycle Research, Progress in Cell Cycle Research. Springer US, Boston, MA, pp. 233–240. 10.1007/978-1-4615-5371-7_18

Feine, O., Hukasova, E., Bruinsma, W., Freire, R., Fainsod, A., Gannon, J., Mahbubani, H.M., Lindqvist, A., Brandeis, M., 2014. Phosphorylation-mediated stabilization of Bora in mitosis coordinates Plx1/Plk1 and Cdk1 oscillations. Cell Cycle Georget. Tex 13, 1727–1736. 10.4161/cc.28630

Friedel, A.M., Pike, B.L., Gasser, S.M., 2009. ATR/Mec1: coordinating fork stability and repair. Curr. Opin. Cell Biol., Cell regulation 21, 237–244. 10.1016/j.ceb.2009.01.017

Fu, Z., Malureanu, L., Huang, J., Wang, W., Li, H., van Deursen, J.M., Tindall, D.J., Chen, J., 2008. Plk1-dependent phosphorylation of FoxM1 regulates a transcriptional programme required for mitotic progression. Nat. Cell Biol. 10, 1076–1082. 10.1038/ncb1767

Gheghiani, L., Loew, D., Lombard, B., Mansfeld, J., Gavet, O., 2017. PLK1 Activation in Late G2 Sets Up Commitment to Mitosis. Cell Rep. 19, 2060–2073. 10.1016/j.celrep.2017.05.031

Grallert, A., Boke, E., Hagting, A., Hodgson, B., Connolly, Y., Griffiths, J.R., Smith, D.L., Pines, J., Hagan, I.M., 2015. A PP1-PP2A phosphatase relay controls mitotic progression. Nature 517, 94–98. 10.1038/nature14019

Hoffmann, I., Clarke, P.R., Marcote, M.J., Karsenti, E., Draetta, G., 1993. Phosphorylation and activation of human cdc25-C by cdc2--cyclin B and its involvement in the self-amplification of MPF at mitosis. EMBO J. 12, 53–63. 10.1002/j.1460-2075.1993.tb05631.x

Hukasova, E., Silva Cascales, H., Kumar, S.R., Lindqvist, A., 2012. Monitoring kinase and phosphatase activities through the cell cycle by ratiometric FRET. J. Vis. Exp. JoVE e3410. 10.3791/3410

Kong, A., Mehanna, H., 2021. WEE1 Inhibitor: Clinical Development. Curr. Oncol. Rep. 23, 107. 10.1007/s11912-021-01098-8

Kumagai, A., Dunphy, W.G., 1992. Regulation of the cdc25 protein during the cell cycle in Xenopus extracts. Cell 70, 139–151. 10.1016/0092-8674(92)90540-S

Kumagai, A., Dunphy, W.G., 1991. The cdc25 protein controls tyrosine dephosphorylation of the cdc2 protein in a cell-free system. Cell 64, 903–914. 10.1016/0092-8674(91)90315-p

Laoukili, J., Alvarez, M., Meijer, L.A.T., Stahl, M., Mohammed, S., Kleij, L., Heck, A.J.R., Medema, R.H., 2008. Activation of FoxM1 during G2 requires cyclin A/Cdk-dependent relief of autorepression by the FoxM1 N-terminal domain. Mol. Cell. Biol. 28, 3076–3087. 10.1128/MCB.01710-07

Lee, J.W., Parameswaran, J., Sandoval-Schaefer, T., Eoh, K.J., Yang, D.-H., Zhu, F., Mehra, R., Sharma, R., Gaffney, S.G., Perry, E.B., Townsend, J.P., Serebriiskii, I.G., Golemis, E.A., Issaeva, N., Yarbrough, W.G., Koo, J.S., Burtness, B., 2019. Combined Aurora Kinase A (AURKA) and WEE1 Inhibition Demonstrates Synergistic Antitumor Effect in Squamous Cell Carcinoma of the Head and Neck. Clin. Cancer Res. Off. J. Am. Assoc. Cancer Res. 25, 3430–3442. 1078-0432.CCR-18-0440

Lemmens, B., Hegarat, N., Akopyan, K., Sala-Gaston, J., Bartek, J., Hochegger, H., Lindqvist, A., 2018. DNA Replication Determines Timing of Mitosis by Restricting CDK1 and PLK1 Activation. Mol. Cell 71, 117-128.e3. 10.1016/j.molcel.2018.05.026

Lemmens, B., Lindqvist, A., 2019. DNA replication and mitotic entry: A brake model for cell cycle progression. J. Cell Biol. 218, 3892–3902. 10.1083/jcb.201909032

Lindqvist, A., Rodríguez-Bravo, V., Medema, R.H., 2009. The decision to enter mitosis: feedback and redundancy in the mitotic entry network. J. Cell Biol. 185, 193–202. 10.1083/jcb.200812045

Lindqvist, A., van Zon, W., Karlsson Rosenthal, C., Wolthuis, R.M.F., 2007. Cyclin B1-Cdk1 activation continues after centrosome separation to control mitotic progression. PLoS Biol. 5, e123. 10.1371/journal.pbio.0050123

Liu, F., Stanton, J.J., Wu, Z., Piwnica-Worms, H., 1997. The human Myt1 kinase preferentially phosphorylates Cdc2 on threonine 14 and localizes to the endoplasmic reticulum and Golgi complex. Mol. Cell. Biol. 17, 571–583. 10.1128/mcb.17.2.571

Lobjois, V., Froment, C., Braud, E., Grimal, F., Burlet-Schiltz, O., Ducommun, B., Bouche, J.-P., 2011. Study of the docking-dependent PLK1 phosphorylation of the CDC25B phosphatase. Biochem. Biophys. Res. Commun. 410, 87–90. 10.1016/j.bbrc.2011.05.110

Lockhead, S., Moskaleva, A., Kamenz, J., Chen, Y., Kang, M., Reddy, A.R., Santos, S.D.M., Ferrell, J.E., 2020. The Apparent Requirement for Protein Synthesis during G2 Phase Is due to Checkpoint Activation. Cell Rep. 32, 107901. 10.1016/j.celrep.2020.107901

Lüscher-Firzlaff, J.M., Lilischkis, R., Lüscher, B., 2006. Regulation of the transcription factor FOXM1c by Cyclin E/CDK2. FEBS Lett. 580, 1716–1722. 10.1016/j.febslet.2006.02.021

Macůrek, L., Lindqvist, A., Lim, D., Lampson, M.A., Klompmaker, R., Freire, R., Clouin, C., Taylor, S.S., Yaffe, M.B., Medema, R.H., 2008. Polo-like kinase-1 is activated by aurora A to promote checkpoint recovery. Nature 455, 119–123. 10.1038/nature07185

Mailand, N., Bekker-Jensen, S., Bartek, J., Lukas, J., 2006. Destruction of Claspin by SCFβTrCP Restrains Chk1 Activation and Facilitates Recovery from Genotoxic Stress. Mol. Cell 23, 307–318. 10.1016/j.molcel.2006.06.016

Mailand, N., Falck, J., Lukas, C., Syljuâsen, R.G., Welcker, M., Bartek, J., Lukas, J., 2000. Rapid destruction of human Cdc25A in response to DNA damage. Science 288, 1425–1429. 10.1126/science.288.5470.1425

Major, M.L., Lepe, R., Costa, R.H., 2004. Forkhead box M1B transcriptional activity requires binding of Cdk-cyclin complexes for phosphorylation-dependent recruitment of p300/CBP coactivators. Mol. Cell. Biol. 24, 2649–2661. 10.1128/mcb.24.7.2649-2661.2004

Mamely, I., van Vugt, M.A., Smits, V.A., Semple, J.I., Lemmens, B., Perrakis, A., Medema, R.H., Freire, R., 2006. Polo-like Kinase-1 Controls Proteasome-Dependent Degradation of Claspin during Checkpoint Recovery. Curr. Biol. 16, 1950–1955. 10.1016/j.cub.2006.08.026

Michael, W.M., Ott, R., Fanning, E., Newport, J., 2000. Activation of the DNA Replication Checkpoint Through RNA Synthesis by Primase. Science 289, 2133–2137. 10.1126/science.289.5487.2133

Minshull, J., Blow, J.J., Hunt, T., 1989. Translation of cyclin mRNA is necessary for extracts of activated Xenopus eggs to enter mitosis. Cell 56, 947–956. 10.1016/0092-8674(89)90628-4

Murakami, H., Aiba, H., Nakanishi, M., Murakami-Tonami, Y., 2010. Regulation of yeast forkhead transcription factors and FoxM1 by cyclin-dependent and polo-like kinases. Cell Cycle 9, 3253–3262. 10.4161/cc.9.16.12599

Nakajima, H., Toyoshima-Morimoto, F., Taniguchi, E., Nishida, E., 2003. Identification of a Consensus Motif for Plk (Polo-like Kinase) Phosphorylation Reveals Myt1 as a Plk1 Substrate*. J. Biol. Chem. 278, 25277–25280. 10.1074/jbc.C300126200

Okumura, E., Morita, A., Wakai, M., Mochida, S., Hara, M., Kishimoto, T., 2014. Cyclin B-Cdk1 inhibits protein phosphatase PP2A-B55 via a Greatwall kinase-independent mechanism. J. Cell Biol. 204, 881–889. 10.1083/jcb.201307160

Parrilla, A., Cirillo, L., Thomas, Y., Gotta, M., Pintard, L., Santamaria, A., 2016. Mitotic entry: The interplay between Cdk1, Plk1 and Bora. Cell Cycle Georget. Tex 15, 3177–3182. 10.1080/15384101.2016.1249544

Peschiaroli, A., Dorrello, N.V., Guardavaccaro, D., Venere, M., Halazonetis, T., Sherman, N.E., Pagano, M., 2006. SCFβTrCP-Mediated Degradation of Claspin Regulates Recovery from the DNA Replication Checkpoint Response. Mol. Cell 23, 319–329. 10.1016/j.molcel.2006.06.013

Pines, J., Hunter, T., 1989. Isolation of a human cyclin cDNA: evidence for cyclin mRNA and protein regulation in the cell cycle and for interaction with p34cdc2. Cell 58, 833–846. 10.1016/0092-8674(89)90936-7

Pomerening, J.R., Sontag, E.D., Ferrell, J.E., 2003. Building a cell cycle oscillator: hysteresis and bistability in the activation of Cdc2. Nat. Cell Biol. 5, 346–351. 10.1038/ncb954

Rata, S., Suarez Peredo Rodriguez, M.F., Joseph, S., Peter, N., Echegaray Iturra, F., Yang, F., Madzvamuse, A., Ruppert, J.G., Samejima, K., Platani, M., Alvarez-Fernandez, M., Malumbres, M., Earnshaw, W.C., Novak, B., Hochegger, H., 2018. Two Interlinked Bistable Switches Govern Mitotic Control in Mammalian Cells. Curr. Biol. CB 28, 3824-3832.e6. 10.1016/j.cub.2018.09.059

Reinhardt, H.C., Aslanian, A.S., Lees, J.A., Yaffe, M.B., 2007. p53-deficient cells rely on ATM-and ATR-mediated checkpoint signaling through the p38MAPK/MK2 pathway for survival after DNA damage. Cancer Cell 11, 175–189. 10.1016/j.ccr.2006.11.024

Roshak, A.K., Capper, E.A., Imburgia, C., Fornwald, J., Scott, G., Marshall, L.A., 2000. The human polo-like kinase, PLK, regulates cdc2/cyclin B through phosphorylation and activation of the cdc25C phosphatase. Cell. Signal. 12, 405–411. 10.1016/S0898-6568(00)00080-2

Saldivar, J.C., Hamperl, S., Bocek, M.J., Chung, M., Bass, T.E., Cisneros-Soberanis, F., Samejima, K., Xie, L., Paulson, J.R., Earnshaw, W.C., Cortez, D., Meyer, T., Cimprich, K.A., 2018. An intrinsic S/G2 checkpoint enforced by ATR. Science 361, 806–810. 10.1126/science.aap9346

Seki, A., Coppinger, J.A., Jang, C.-Y., Yates, J.R., Fang, G., 2008. Bora and Aurora A Cooperatively Activate Plk1 and Control the Entry into Mitosis. Science 320, 1655–1658. 10.1126/science.1157425

Serpico, A.F., D’Alterio, G., Vetrei, C., Della Monica, R., Nardella, L., Visconti, R., Grieco, D., 2019. Wee1 Rather Than Plk1 Is Inhibited by AZD1775 at Therapeutically Relevant Concentrations. Cancers 11, 819. 10.3390/cancers11060819

Sha, W., Moore, J., Chen, K., Lassaletta, A.D., Yi, C.-S., Tyson, J.J., Sible, J.C., 2003. Hysteresis drives cell-cycle transitions in Xenopus laevis egg extracts. Proc. Natl. Acad. Sci. 100, 975–980. 10.1073/pnas.0235349100

Spencer, S.L., Cappell, S.D., Tsai, F.-C., Overton, K.W., Wang, C.L., Meyer, T., 2013. The proliferation-quiescence decision is controlled by a bifurcation in CDK2 activity at mitotic exit. Cell 155, 369–383. 10.1016/j.cell.2013.08.062

Steegmaier, M., Hoffmann, M., Baum, A., Lénárt, P., Petronczki, M., Krssák, M., Gürtler, U., Garin-Chesa, P., Lieb, S., Quant, J., Grauert, M., Adolf, G.R., Kraut, N., Peters, J.-M., Rettig, W.J., 2007. BI 2536, a potent and selective inhibitor of polo-like kinase 1, inhibits tumor growth in vivo. Curr. Biol. CB 17, 316–322. 10.1016/j.cub.2006.12.037

Sur, S., Agrawal, D.K., 2016. Phosphatases and kinases regulating CDC25 activity in the cell cycle: clinical implications of CDC25 overexpression and potential treatment strategies. Mol. Cell. Biochem. 416, 33–46. 10.1007/s11010-016-2693-2

Tavernier, N., Noatynska, A., Panbianco, C., Martino, L., Van Hove, L., Schwager, F., Léger, T., Gotta, M., Pintard, L., 2015. Cdk1 phosphorylates SPAT-1/Bora to trigger PLK-1 activation and drive mitotic entry in C. elegans embryos. J. Cell Biol. 208, 661–669. 10.1083/jcb.201408064

Thomas, Y., Cirillo, L., Panbianco, C., Martino, L., Tavernier, N., Schwager, F., Van Hove, L., Joly, N., Santamaria, A., Pintard, L., Gotta, M., 2016. Cdk1 Phosphorylates SPAT-1/Bora to Promote Plk1 Activation in C. elegans and Human Cells. Cell Rep. 15, 510–518. 10.1016/j.celrep.2016.03.049

Toledo, L.I., Altmeyer, M., Rask, M.-B., Lukas, C., Larsen, D.H., Povlsen, L.K., Bekker-Jensen, S., Mailand, N., Bartek, J., Lukas, J., 2013. ATR Prohibits Replication Catastrophe by Preventing Global Exhaustion of RPA. Cell 155, 1088–1103. 10.1016/j.cell.2013.10.043

Vigneron, S., Sundermann, L., Labbé, J.-C., Pintard, L., Radulescu, O., Castro, A., Lorca, T., 2018. Cyclin A-cdk1-Dependent Phosphorylation of Bora Is the Triggering Factor Promoting Mitotic Entry. Dev. Cell 45, 637-650.e7. 10.1016/j.devcel.2018.05.005

Wang, Z., Li, W., Li, F., Xiao, R., 2024. An update of predictive biomarkers related to WEE1 inhibition in cancer therapy. J. Cancer Res. Clin. Oncol. 150, 13. 10.1007/s00432-023-05527-y

Watanabe, N., Arai, H., Iwasaki, J.-I., Shiina, M., Ogata, K., Hunter, T., Osada, H., 2005. Cyclin-dependent kinase (CDK) phosphorylation destabilizes somatic Wee1 via multiple pathways. Proc. Natl. Acad. Sci. U. S. A. 102, 11663–11668. 10.1073/pnas.0500410102

Wierstra, I., Alves, J., 2006. FOXM1c is activated by cyclin E/Cdk2, cyclin A/Cdk2, and cyclin A/Cdk1, but repressed by GSK-3alpha. Biochem. Biophys. Res. Commun. 348, 99–108. 10.1016/j.bbrc.2006.07.008

Wright, G., Golubeva, V., Remsing Rix, L.L., Berndt, N., Luo, Y., Ward, G.A., Gray, J.E., Schonbrunn, E., Lawrence, H.R., Monteiro, A.N.A., Rix, U., 2017. Dual Targeting of WEE1 and PLK1 by AZD1775 Elicits Single Agent Cellular Anticancer Activity. ACS Chem. Biol. 12, 1883–1892. 10.1021/acschembio.7b00147

Zhang, J., Yuan, C., Wu, J., Elsayed, Z., Fu, Z., 2015. Polo-like kinase 1-mediated phosphorylation of Forkhead box protein M1b antagonizes its SUMOylation and facilitates its mitotic function. J. Biol. Chem. 290, 3708–3719. 10.1074/jbc.M114.634386

