## Supplementary data for "Wee1 inhibition decouples Cdk1 and Plk1 activities: a role for gradual Cdk1 activation throughout G2 phase"

**Figure S1. Duration of mitosis after adding Wee1i to hTERT RPE1 Cyclin B1-YFP cells.**

**A**. Quantification of Cyclin B1-YFP fluorescence of hTERT RPE1 Cyclin B1-YFP cells *in-silico* synchronized on mitotic entry.

**B**. Average signal and standard deviation from A.

**C, D.** RPE1 Cyclin B1-YFP cells were monitored by time-lapse microscopy upon addition of MK1775. C, duration of mitosis (x-axis) is plotted versus Cyclin B1-YFP level at mitotic entry (y-axis) D, duration of mitosis (y-axis) is plotted versus estimated time before mitosis should Wee1 inhibitors not have been added (x-axis). The estimate is based on Cyclin B1-YFP accumulation of control cells in B.

**Figure S2. Trendline used to estimate position within G2 phase for Figure 4.**

U2OS Cyclin B1-YFP cells were monitored by live cell microscopy. Plots show single cell traces (left) and average (right) of YFP fluorescence after *in silico* synchronization in mitosis. Cell cycle position for individual cells in Figure 4E was estimated by comparison to the average trend line.

### Figure S1

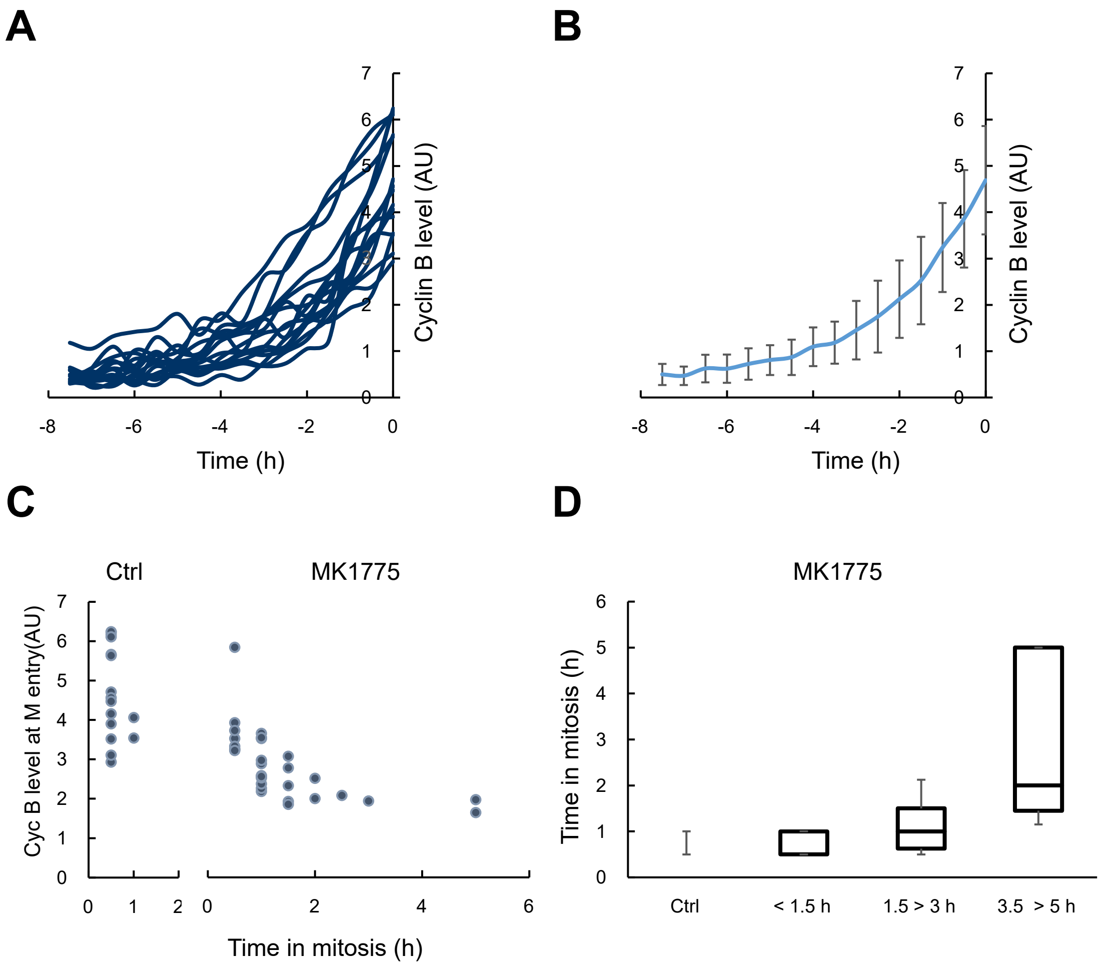
A
B
C
D
Cyclin B level (AU)
Time (h)
Ctrl
MK1775
Cyc B level at M entry(AU)
Time in mitosis (h)
< 1.5 h
1.5 > 3 h
3.5 > 5 h

### Figure S2

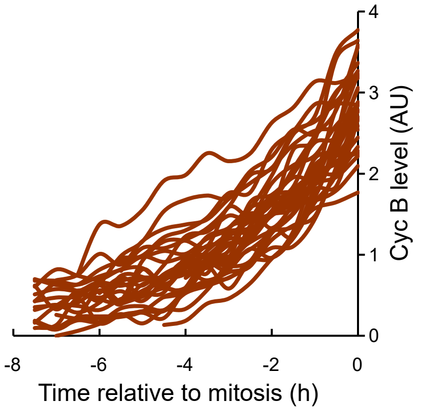

Cyc B level (AU)
0
1
2
3
4
-8
-6
-4
-2
0
Time relative to mitosis (h)


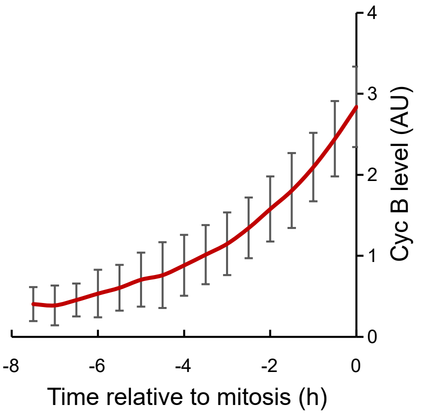

Cyc B level (AU)
0
1
2
3
4
-8
-6
-4
-2
0
Time relative to mitosis (h)

### Appendix: cell cycle model

The model is based on ordinary differential equations. The aim has been to limit the number of parameters to avoid overparameterization and to allow fitting to experimental data.

All equations in the model are assigned into 3 groups corresponding to cell cycle phases in which they are considered most relevant. The following labels were added to protein names to indicate: “T” – total level, “P” – phosphorylated, “A” – active, “I” – inactive.

For some proteins the total level is defined as constant = 1.

There are two parameters in the model that direct protein synthesis:

1. *Slow constitutive synthesis* is a constant (specified as E2F in Fig. 3). The constant is different for every protein.
2. *Active synthesis* denotes regulated expression. For simplicity, factors are given as the species FoxM (specified as FoxM/NF-Y in Fig. 3).

#### G2 phase

##### FoxM

The level of active FoxM [$FoxM_A$] is regulated by cell cycle kinases Cdk1, Cdk2(Laoukili et al., 2008; Lüscher-Firzlaff et al., 2006; Major et al., 2004; Saldivar et al., 2018) and Plk1(Fu et al., 2008, p. 1; Murakami et al., 2010; Zhang et al., 2015).

$$\frac{d[FoxM_A]}{dt} = \frac{k_{cycbfoxm} \cdot [CycB_A] \cdot [FoxM_I]}{1 + [FoxM_I]} + \frac{k_{cycafoxm} \cdot [CycA_A] \cdot [FoxM_I]}{1 + [FoxM_I]} + \frac{k_{cycefoxm} \cdot [CycE_A] \cdot [FoxM_I]}{1 + [FoxM_I]} + \frac{k_{plk1foxm} \cdot [Plk1_A] \cdot [FoxM_I]}{1 + [FoxM_I]} - \frac{k_{pp2foxm} \cdot [PP2_A] \cdot [FoxM_A]}{1 + [FoxM_A]}$$

$$[FoxM_I] = [FoxM_T] - [FoxM_A]$$

##### CycBCdk1

The term CycBCdk1 denotes both Cyclin B-Cdk1 and Cyclin A-Cdk1 complexes. The total level of CycBCdk1 is described by the following equation:

$$\frac{d[CycB_T]}{dt} = k_{scycb} + V_{scycbfoxm} - V_{dcycb} \cdot [CycB_T]$$

Where $k_{scycb}$ is a constant corresponding to slow constitutive synthesis, $V_{scycbfoxm}$ is a rate function describing the regulation of *Active synthesis*. And $V_{dcycb}$ function describes the degradation. Here, we assume that all Cyclin B binds immediately to Cdk1 forming Cyclin B-Cdk1 complexes.

The level of active Cyclin B-Cdk1 [$CycB_A$] is regulated by the balance between Cdc25C phosphatase and Wee1/Myt1 kinases. The inhibitory phosphorylation on Thr 14 and Tyr 15 by Wee1/Myt1 (Fattaey and Booher, 1997; Liu et al., 1997, p. 1) turns Cdk1 to an inactive phosphorylated form [$CycB_I$] and Cdc25 turns it back to an active form by removing those phosphogroups (Hoffmann et al., 1993; Kumagai and Dunphy, 1992, 1991).

$$\frac{d[CycB_A]}{dt} = \frac{k_{cdc25cycb} \cdot [Cdc25C_A] \cdot [CycB_I]}{1 + [CycB_I]} + \frac{k_{cdc25abcycb} \cdot [Cdc25AB_A] \cdot [CycB_I]}{1 + [CycB_I]} - \frac{k_{wee1cycb} \cdot [Wee1_A] \cdot [CycB_A]}{1 + [CycB_A]}$$

The level of inactive Cyclin B-Cdk1 [$CycB_I$] is defined by:

$$[CycB_I] = [CycB_T] - [CycB_A]$$

**CycACdk2**

The term CycACdk2 denotes Cyclin A-Cdk2 complexes. The total level of CycACdk2 is given by:

$$\frac{d[CycA_T]}{dt} = k_{scyca} + V_{scycafoxm} - V_{dcyca} \cdot [CycA_T]$$

Where $k_{scyca}$ is a constant corresponding to synthesis, $V_{scycafoxm}$ is a rate function describing the regulation of *Active synthesis* and $V_{dcyca}$ function describes the degradation of Cyclin A.

Inhibition/Association of CycA by p27:

$$\frac{d[CycAp27]}{dt} = k_{assa} \cdot [CycA_A] \cdot ([p27_T] - [CycAp27] - [CycEp27]) - \left(k_{disa} + V_{dp27} + V_{dcyca}\right) \cdot [CycAp27]$$

The level of active CycA-Cdk2 is regulated by Cdc25AB and Wee1:

$$\frac{d[CycA_A]}{dt} = \frac{k_{cdc25cyca} \cdot [Cdc25AB_A] \cdot [CycA_I]}{1 + [CycA_I]} - \frac{k_{wee1cyca} \cdot [Wee1_A] \cdot [CycA_A]}{1 + [CycA_A]}$$

The level of inactive CycA-Cdk2 is defined by:

$$[CycA_I] = [CycA_T] - [CycAp27] - [CycA_A]$$

**Plk1**

The total level of Plk1 is defined by:

$$\frac{d[Plk1_T]}{dt} = k_{splk1} + V_{splk1foxm} - V_{dplk1} \cdot [Plk1_T]$$

$k_{splk1}$ is a constant corresponding to S*low constitutive synthesis*, $V_{splk1foxm}$ describes the regulation of *Active synthesis*, and $V_{dplk1}$ function describes the degradation.

The level of active Plk1 is regulated by Aurora A kinase with cofactor Bora [AurBora] (Macůrek et al., 2008; Seki et al., 2008) and Chk1 (Bruinsma et al., 2017):

$$\frac{d[Plk1_A]}{dt} = \frac{k_{aurboraplk1} \cdot [AurBora] \cdot [Plk1_I]}{1 + [Plk1_I]} - \frac{k_{chk1plk1} \cdot [Chk1_A] \cdot [Plk1_A]}{1 + [Plk1_A]}$$

The level of inactive Plk1 is defined by:

$$[Plk1_I] = [Plk1_T] - [Plk1_A]$$

**Aurora A**

The total level of Aurora A is defined by:

$$\frac{d[AurA_T]}{dt} = k_{saura} + V_{saurafoxm} - V_{daur} \cdot [AurA_T]$$

$k_{saura}$ is a constant corresponding to S*low constitutive synthesis*, $V_{saurafoxm}$ describes the regulation of *Active synthesis*, and $V_{daur}$ function describes the degradation.

For simplicity we do not involve an Aurora activation – inactivation path. We assume that Aurora A is always active:

$$[AurA] = [AurA_T] - [AurBora]$$

**PP2**

*We use [PP2] as the collective value of all phosphatases opposing Cdk1* (Grallert et al., 2015).

Cdk1 is involved in inactivation of PP2 (by Greatwall) (Okumura et al., 2014). For activation of PP2 we use the constant ($k_{pp2pp2}$)

$$\frac{d[PP2_I]}{dt} = \frac{k_{cycbpp2} \cdot [CycB_A] \cdot [PP2_A]}{1 + [PP2_A]} - \frac{k_{pp2pp2} \cdot [PP2_I]}{1 + [PP2_I]}$$

The level of active PP2 is defined by:

$$[PP2_A] = [PP2_T] - [PP2_I]$$

**Bora**

In our model Bora is present in 2 forms: not phosphorylated by Cdk1 [Bora] and phosphorylated by Cdk1 [$Bora_P$](Thomas et al., 2016; Vigneron et al., 2018).

Plk1 is involved in the degradation of Bora [Bora] while Cdk1 and Cdk2 can phosphorylate Bora which leads to stabilization (Feine et al., 2014).

$$\frac{d[Bora]}{dt} = k_{sbora} + V_{sborafoxm} + \frac{k_{pp2bora} \cdot [PP2_A] \cdot [Bora_P]}{1 + [Bora_P]} - \frac{k_{cycbbora} \cdot [CycB_A] \cdot [Bora]}{1 + [Bora]} - V_{dbora} \cdot [Bora] - V_{dboraplk1} \cdot [Bora]$$

$$\frac{d[Bora_P]}{dt} = \frac{k_{cycbbora} \cdot [CycB_A] \cdot [Bora]}{1 + [Bora]} + \frac{k_{cycabora} \cdot [CycA_A] \cdot [Bora]}{1 + [Bora]} - \frac{k_{pp2bora} \cdot [PP2_A] \cdot [Bora_P]}{1 + [Bora_P]} - V_{dbora} \cdot [Bora_P]$$

**Aurora A - Bora**

Aurora A form a complex with Bora phosphorylated by Cdk1 [AurBora] (Parrilla et al., 2016).

$$\frac{d[AurBora]}{dt} = k_{assb} \cdot [AurA] \cdot [Bora_P] - (k_{disb} + V_{daur} + V_{dbora}) \cdot [AurBora]$$

**Cdc25C**

The total level of Cdc25C is defined as a constant.

For activation of Cdc25C we use the following equation:

$$\frac{d[Cdc25C_A]}{dt} = \frac{k_{cycbcdc25} \cdot [CycB_A] \cdot [Cdc25C_I]}{1 + [Cdc25C_I]} + \frac{k_{plk1cdc25} \cdot [Plk1_A] \cdot [Cdc25C_I]}{1 + [Cdc25C_I]} + \frac{k_{aurcdc25} \cdot [AurA] \cdot [Cdc25C_I]}{1 + [Cdc25C_I]} - \frac{k_{pp2cdc25} \cdot [PP2_A] \cdot [Cdc25C_A]}{1 + [Cdc25C_A]} - \frac{k_{chk1cdc25} \cdot [Chk1_A] \cdot [Cdc25C_A]}{1 + [Cdc25C_A]}$$

Cdk1 as well as Plk1 and Aurora A promote activation (Gheghiani et al., 2017; Roshak et al., 2000; Sur and Agrawal, 2016), and PP2 and Chk1 promote inactivation of Cdc25C (Sur and Agrawal, 2016).

The level of inactive Cdc25C is defined by:

$$[Cdc25C_I] = [Cdc25C_T] - [Cdc25C_A]$$

**Wee1**

The total level of Wee1 is defined as a constant.

Inactivation of Wee1 by Cdk and Plk1 (Nakajima et al., 2003; Watanabe et al., 2005):

$$\frac{d[Wee1_I]}{dt} = \frac{k_{cycbwee1} \cdot [CycB_A] \cdot [Wee1_A]}{1 + [Wee1_A]} + \frac{k_{plk1wee1} \cdot [Plk1_A] \cdot [Wee1_A]}{1 + [Wee1_A]} + \frac{k_{cycawee1} \cdot [CycA_A] \cdot [Wee1_A]}{1 + [Wee1_A]} + \frac{k_{cycewee1} \cdot [CycE_A] \cdot [Wee1_A]}{1 + [Wee1_A]} - \frac{k_{pp2wee1} \cdot [PP2_A] \cdot [Wee1_I]}{1 + [Wee1_I]}$$

The level of active Wee1 is defined by:

$$[Wee1_A] = [Wee1_T] - [Wee1_I]$$

### S-phase

**CycECdk2**

$$\frac{d[CycE_T]}{dt} = k_{scyce} - V_{dcyce} \cdot [CycE_T]$$

$$\frac{d[CycEp27]}{dt} = k_{asse} \cdot [CycE_A] \cdot ([p27_T] - [CycAp27] - [CycEp27]) - \left(k_{dise} + V_{dp27} + V_{dcyce}\right) \cdot [CycEp27]$$

$$\frac{d[CycE_A]}{dt} = \frac{k_{cdc25cyce} \cdot [Cdc25AB_A] \cdot [CycE_I]}{1 + [CycE_I]} - \frac{k_{wee1cyce} \cdot [Wee1_A] \cdot [CycE_A]}{1 + [CycE_A]}$$

$$[CycE_I] = [CycE_T] - [CycEp27] - [CycE_A]$$

**Cdc25AB**

For simplicity we combined two isoforms of Cdc25 (Cdc25A and Cdc25B) [Cdc25AB].

The total level of Cdc25AB is defined as a constant.

For activation of Cdc25AB we used the following equation:

$$\frac{d[Cdc25AB_A]}{dt} = \frac{k_{cycecdc25a} \cdot [CycE_A] \cdot [Cdc25AB_I]}{1 + [Cdc25AB_I]} + \frac{k_{cycacdc25a} \cdot [CycA_A] \cdot [Cdc25AB_I]}{1 + [Cdc25AB_I]} + \frac{k_{plk1cdc25} \cdot [Plk1_A] \cdot [Cdc25AB_I]}{1 + [Cdc25AB_I]} + \frac{k_{aurcdc25} \cdot [AurA] \cdot [Cdc25AB_I]}{1 + [Cdc25AB_I]} - \frac{k_{pp2cdc25} \cdot [PP2_A] \cdot [Cdc25AB_A]}{1 + [Cdc25AB_A]} - \frac{k_{chk1cdc25ab} \cdot [Chk1_A] \cdot [Cdc25AB_A]}{1 + [Cdc25AB_A]}$$

Cdk2, Plk1 and Aurora A activate Cdc25AB, whereas PP2 and Chk1 inactivate Cdc25AB (Lobjois et al., 2011; Mailand et al., 2000; Reinhardt et al., 2007; Sur and Agrawal, 2016).

The level of inactive Cdc25AB is defined by:

$$[Cdc25AB_I] = [Cdc25AB_T] - [Cdc25AB_A]$$

**ATR**

The total level of ATR is defined as a constant.

For activation of ATR we used the following equation:

$$\frac{d[Atr_A]}{dt} = \frac{k_{replatr} \cdot [Origin_F :: Replication] \cdot [Atr_I]}{1 + [Atr_I]} - \frac{k_{autatr} \cdot [Atr_A]}{1 + [Atr_A]}$$

ATR is activated by Replication (Friedel et al., 2009; Toledo et al., 2013). For simplicity we used a constant $k_{autatr}$ to denote inactivation of ATR.

The level of inactive ATR is defined by:

$$[Atr_I] = [Atr_T] - [Atr_A]$$

**Chk1**

The total level of ATR is defined as a constant.

ATR-dependent Chk1 activation is described by:

$$\frac{d[Chk1_A]}{dt} = \frac{k_{atrchk1} \cdot [Atr_A] \cdot [Chk1_I]}{1 + [Chk1_I]} - \frac{k_{plk1chk1} \cdot [Plk1_A] \cdot [Chk1_A]}{1 + [Chk1_A]} - \frac{k_{autchk1} \cdot [Chk1_A]}{1 + [Chk1_A]}$$

Plk1 inactivates Chk1 (Mailand et al., 2006; Mamely et al., 2006; Peschiaroli et al., 2006) by inducing degradation of Claspin, which is required for activation of Chk1.

The level of inactive Chk1 is defined by:

$$[Chk1_I] = [Chk1_T] - [Chk1_A]$$

**Cdc6**

The total level of Cdc6 is defined as a constant.

Cdk2 dependent inactivation of Cdc6 (Coulombe et al., 2019; Dutta and Bell, 1997; Michael et al., 2000):

$$\frac{d[Cdc6_I]}{dt} = \frac{k_{cycecdc6} \cdot [CycE_A] \cdot [Cdc6_A]}{1 + [Cdc6_A]} + \frac{k_{cycacdc6} \cdot [CycA_A] \cdot [Cdc6_A]}{1 + [Cdc6_A]}$$

The level of active Cdc6 is defined by:

$$[Cdc6_A] = [Cdc6_T] - [Cdc6_I]$$

**Cdc7**

The total level of Cdc7 is described as:

$$\frac{d[Cdc7_T]}{dt} = k_{scdc7} - V_{dcdc7} \cdot [Cdc7_T]$$

*The* $k_{scdc7}$ is a constant corresponding to *Slow constitutive synthesis*, whereas $V_{dcdc7}$ function describes the degradation.

Cdk2 dependent Cdc7 activation (Coulombe et al., 2019; Dutta and Bell, 1997; Michael et al., 2000) is described by:

$$\frac{d[Cdc7_A]}{dt} = \frac{k_{cycecdc7} \cdot [CycE_A] \cdot [Cdc7_I]}{1 + [Cdc7_I]} + \frac{k_{cycacdc7} \cdot [CycA_A] \cdot [Cdc7_I]}{1 + [Cdc7_I]}$$

The level of inactive Cdc7 is defined by:

$$[Cdc7_I] = [Cdc7_T] - [Cdc7_A]$$

**DNA replication**

Is described by the following equations:

The first step, loading of Orc, Cdc6, and MCM proteins [Cdc6] onto DNA is described as Origin licensing [$Origin_L$] (Bleichert, 2019; Coulombe et al., 2019; Dutta and Bell, 1997; Lemmens et al., 2018; Michael et al., 2000) :

$$\frac{d[Origin_L]}{dt} = k_{assl} \cdot [Cdc6_A] \cdot [Origin] - \frac{k_{assf} \cdot [Cdc7_A] \cdot [Origin_L]}{1 + [Origin_L]} - V_{repl} \cdot [Origin_L]$$

The second step is described as Origin firing by Cdc7 (Coulombe et al., 2019; Dutta and Bell, 1997; Lemmens et al., 2018; Michael et al., 2000):

$$\frac{d[Origin_F :: Replication]}{dt} = \frac{k_{assf} \cdot [Cdc7_A] \cdot [Origin_L]}{1 + [Origin_L]} - [Origin_F :: Replication] \cdot [Origin_D]^{kodr}$$

$$\frac{d[Origin_D]}{dt} = V_{repl} \cdot ([Origin_T] - [Origin_D])$$

$$[Origin] = [Origin_T] - [Origin_L] - [Origin_F :: Replication]$$

#### G1-phase, taken from (Barr et al., 2016).

$$\frac{d[Skp2]}{dt} = k_{sskp2} - \left(k_{dskp2} + k_{dskp2c1} \cdot [Cdh1]\right) \cdot [Skp2]$$

$$\frac{d[Cdh1]}{dt} = k_{acdh1} \cdot (Cdh1_T - [EmiC] - [Cdh1]) - V_{icdh1} \cdot [Cdh1] - k_{asec} \cdot [Cdh1] \cdot ([Emi1_T] - [EmiC]) + (k_{diec} + k_{demi1}) \cdot ([Cdh1dp] - [Cdh1])$$

$$\frac{d[EmiC]}{dt} = k_{asec} \cdot (Cdh1_T - [EmiC]) \cdot ([Emi1_T] - [EmiC]) - (k_{diec} + k_{demi1}) \cdot [EmiC]$$

$$\frac{d[Cdh1dp]}{dt} = k_{acdh1} - ([Cdh1_T] - [Cdh1dp]) - V_{icdh1} \cdot [Cdh1dp]$$

$$\frac{d[p27_T]}{dt} = k_{s27} - V_{dp27} \cdot [p27_T]$$

#### Global Quantities (Type: assignment)

$$V_{icdh1} = k_{icdh1e} \cdot [CycE_A] + k_{icdh1a} \cdot [CycA_A]$$

$$V_{dcyce} = k_{dcyce} + k_{dcycee} \cdot [CycE_A] + k_{dcycea} \cdot [CycA_A]$$

$$V_{dcyca} = k_{dcyca} + k_{dcycac1} \cdot [Cdh1]$$

$$V_{dp27} = (k_{d27e} \cdot [CycE_A] + k_{d27a} \cdot [CycA_A]) \cdot [Skp2] + k_{d27}$$

$$V_{dcycb} = k_{dcycb} + k_{dcycbc1} \cdot [Cdh1]$$

$$V_{dplk1} = k_{dplk1} + k_{dplk1c1} \cdot [Cdh1]$$

$$V_{daur} = k_{daur} + k_{daurc1} \cdot [Cdh1]$$

$$V_{dbora} = k_{dbora} + k_{dborac1} \cdot [Cdh1]$$

$$V_{dboraplk1} = k_{dboraplk1} \cdot [Plk1_A]$$

$$V_{scycbfoxm} = k_{scycbfoxm} \cdot [FoxM_A]$$

$$V_{saurafoxm} = k_{saurafoxm} \cdot [FoxM_A]$$

$$V_{splk1foxm} = k_{splk1foxm} \cdot [FoxM_A]$$

$$V_{scycafoxm} = k_{scycafoxm} \cdot [FoxM_A]$$

$$V_{sborafoxm} = k_{sborafoxm} \cdot [FoxM_A]$$

$$V_{dcdc7} = k_{dcdc7} + k_{dcdc7c1} \cdot [Cdh1]$$

$$V_{repl} = k_{repl} \cdot [Origin_F :: Replication]$$

#### Global Quantities (Type: fixed)

| # | Name | Value | | # | Name | Value | | # | Name | Value |
|---|---|---|---|---|---|---|---|---|---|---|
| 1 | ksskp2 | 0.004 | | 35 | kdaurc1 | 0.75090424 | | 69 | kchk1wee1 | 200 |
| 2 | kdskp2 | 0.002 | | 36 | kcycbcdc25 | 0.101625007 | | 70 | kchk1cdc25 | 10 |
| 3 | kdskp2c1 | 0.2 | | 37 | kpp2cdc25 | 0.09027315 | | 71 | kreplatr | 0.5 |
| 4 | Cdh1T | 1 | | 38 | kplk1cdc25 | 2.09183084 | | 72 | katrchk1 | 0.3 |
| 5 | kacdh1 | 0.02 | | 39 | kaurcdc25 | 0.1 | | 73 | kautatr | 0.25 |
| 6 | kicdh1e | 0.07 | | 40 | kwee1cycb | 44.72172096 | | 74 | kautchk1 | 0.4 |
| 7 | kicdh1a | 0.2 | | 41 | kcdc25cycb | 4.5289313 | | 75 | kcdc25cyca | 50 |
| 8 | kasec | 2 | | 42 | kautplk1 | 0.05 | | 76 | kcdc25cyce | 50 |
| 9 | kdiec | 0.02 | | 43 | kaurboraplk1 | 0.11 | | 77 | kwee1cyca | 0.5 |
| 10 | ksemi1 | 0.003 | | 44 | kcycbpp2 | 0.155115167 | | 78 | kwee1cyce | 7 |
| 11 | kdemi1 | 0.001 | | 45 | kpp2pp2 | 0.05 | | 79 | kcycacdc25a | 0.1 |
| 12 | ks27 | 0.008 | | 46 | kcycbwee1 | 0.64369118 | | 80 | kcycecdc25a | 0.05 |
| 13 | kd27 | 0.004 | | 47 | kpp2wee1 | 0.658411743 | | 81 | kcycabora | 0.055959715 |
| 14 | kd27e | 2 | | 48 | kplk1wee1 | 0.735261119 | | 82 | kplk1chk1 | 0.5 |
| 15 | kd27a | 2 | | 49 | kcycbprobe | 2 | | 83 | kchk1plk1 | 25 |
| 16 | kscyca | 0.00253 | | 50 | kpp2probe | 1 | | 84 | kscdc7 | 1 |
| 17 | kdcyca | 0.002 | | 51 | kassb | 0.550645521 | | 85 | kdcdc7 | 0.001 |
| 18 | kdcycac1 | 0.4 | | 52 | kdisb | 0.02 | | 86 | kdcdc7c1 | 100 |
| 19 | kassa | 1 | | 53 | kdbora | 0.072778314 | | 87 | kcdc25abcycb | 1.5 |
| 20 | kdisa | 0.02 | | 54 | kdborac1 | 0.567370959 | | 88 | kchk1cdc25ab | 0.1 |
| 21 | kscyce | 0.005 | | 55 | ksbora | 0.002748359 | | 89 | kcycawee1 | 0.05 |
| 22 | kdcyce | 0.001 | | 56 | kpp2bora | 0.096448508 | | 90 | kcycewee1 | 0.8 |
| 23 | kdcycee | 0.0001 | | 57 | kcycbbora | 3.445732371 | | 91 | kodr | 400 |
| 24 | kdcycea | 0.06 | | 58 | kdboraplk1 | 0.011541844 | | 92 | kcycbfoxm | 1.65 |
| 25 | kasse | 1 | | 59 | kcycacdc6 | 5 | | 93 | kcycafoxm | 0.16 |
| 26 | kdise | 0.02 | | 60 | kcycecdc6 | 0.1 | | 94 | kcycefoxm | 0.13 |
| 27 | kscycb | 0.0002 | | 61 | kcycacdc7 | 0.005 | | 95 | kplk1foxm | 0.06 |
| 28 | kdcycb | 0.002 | | 62 | kcycecdc7 | 0.001 | | 96 | kpp2foxm | 1 |
| 29 | kdcycbc1 | 0.35 | | 63 | kassl | 0.1 | | 97 | kscycbfoxm | 0.05 |
| 30 | ksplk1 | 0.003736186 | | 64 | kdisl | 0 | | 98 | ksplk1foxm | 0.135 |
| 31 | kdplk1 | 0.000229651 | | 65 | kassf | 1 | | 99 | kscycafoxm | 0.005 |
| 32 | kdplk1c1 | 0.109550453 | | 66 | kdisf | 0 | | 100 | ksborafoxm | 0.0053 |
| 33 | ksaura | 0.002134667 | | 67 | krepl | 0.0235 | | | | |
| 34 | kdaur | 0.005 | | 68 | ksaurafoxm | 0.049820691 | | | | |
